# Probabilistic Colocalization of Genetic Variants from Complex and Molecular Traits: Promise and Limitations

**DOI:** 10.1101/2020.07.01.182097

**Authors:** Abhay Hukku, Milton Pividori, Francesca Luca, Roger Pique-Regi, Hae Kyung Im, Xiaoquan Wen

## Abstract

Colocalization analysis has emerged as a powerful tool to uncover the overlapping of causal variants responsible for both molecular and complex disease phenotypes. The findings from colocalization analysis yield insights into the molecular pathways of complex diseases. In this paper, we conduct an in-depth investigation of the promise and limitations of the available colocalization analysis approaches. Focusing on variant-level colocalization approaches, we first establish the connections between various existing methods. We proceed to discuss the impacts of various controllable analytical factors and uncontrollable practical factors on outcomes of colocalization analysis through realistic simulations and real data examples. We identify a single analytical factor, the specification of prior enrichment levels, which can lead to severe inflation of false-positive colocalization findings. Meanwhile, the combination of many other analytical and practical factors all lead to diminished power. Consequently, we recommend the following strategies for the best practice of colocalization analysis: i) estimating prior enrichment level from the observed data; and ii) separating fine-mapping and colocalization analysis. Our analysis of 4,091 complex traits and the multi-tissue eQTL data from the GTEx (version 8) suggests that colocalizations of molecular QTLs and GWAS traits are widespread in many complex traits. However, only a small proportion can be confidently identified from currently available data due to a lack of power. Our findings should serve as an important benchmark for the current and future integrative genetic association analysis applications.

## 1 Introduction

The advancements in genetic association analysis of complex and molecular traits have uncovered a large volume of putative causal genetic variants. Subsequently, utilizing the genetic association discoveries to explore the molecular mechanisms of complex disease etiology has become a standard practice in human genetics research. Various types of analytical approaches designed for the integrative analysis of data from expression quantitative trait loci (eQTL) mapping and genome-wide association analysis (GWAS) of complex traits have shown promise in implicating molecular pathways connecting genetic variations, molecular phenotype changes, and complex diseases [1, 2, 3, 4, 5, 6].

Colocalization analysis is one of the integrative analysis techniques that aims to identify a single genetic variant introducing simultaneous phenotypic changes in multiple molecular and/or complex traits. It is closely related to the approaches designed to uncover single-point univariate pleiotropic effects [7], but additionally emphasizes the causality of the target variants (i.e., the observed effect on any trait is unlikely due to linkage disequilibrium). Although colocalization analysis is not constrained by the types of phenotypes investigated, we focus our discussions in this paper on a single complex trait and one type of molecular trait, e.g., gene expressions. The discoveries from this class of analyses have resulted in molecular insights of complex diseases, e.g., atherosclerosis [8], the age of onset of menarche and menopause [9], and cardiovascular disease [10]. Nevertheless, our main conclusions should extend to colocalization analyses of arbitrary traits.

There are two primary types of colocalization analysis approaches in the current literature. The first kind, represented by Regulatory Trait Concordance (RTC) [2] and Joint Likelihood Mapping (JLIM) [11], makes claims that causal GWAS hits and eQTL signals co-exist within a genomic region consisting of tightly linked genetic variants. We refer to this type as the *locus-level colocalization analysis*. The second kind, illustrated by *coloc* [12], *eCAVIAR* [13], and ENLOC/fastENLOC [14, 6], attempts to uncover colocalization signals at the single SNP resolution. We refer to this type as *SNP-level colocalization analysis*. The common obstacle for both types of colocalization analysis is linkage disequilibrium (LD) among candidate SNPs. Hypothetically, with complete linkage equilibrium, colocalization analysis becomes relatively trivial, and various approaches from both types converge. As demonstrated in [14] (see the example of two perfectly linked SNPs), with the presence of a high-level of LD, SNP-level colocalization may not be identifiable from the observed association data, i.e., the likelihood information alone is insufficient for identifying SNP-level colocalization. Consequently, the second type of approaches are all Bayesian approaches and rely on prior information to provide probabilistic quantifications of SNP-level colocalizations. Moreover, SNP-level colocalization analysis becomes synonymous with *probabilistic colocalization analysis*. We provide a more detailed overview and summary regarding both locus- and SNP-level colocalization analysis methods in the Method section, and the main focus of this paper is on the SNP-level colocalization methods.

Probabilistic colocalization analysis faces many challenges, both analytically and practically. Analytical factors, e.g., prior specifications and model assumptions required for likelihood computation (e.g., consideration of allelic heterogeneity), are known to have drastic impacts on analysis outcomes. Practically, even when the ideal analytical strategies are applied, the power of the colocalization analysis can still be limited by the underlying association data, e.g., the power of marginal association analysis. In this paper, we take a divide-and-conquer strategy to systematically investigate various analytical and practical factors in probabilistic colocalization analyses. Through analytical derivation and numerical experiments, we attempt to isolate each factor and quantify its effects on potential false positive and false negative findings. We seek to identify a set of best analytical strategies that enable robust and powerful probabilistic colocalization analysis. Also, we hope to practically illustrate the natural limitations of the colocalization analysis based on the currently available data and establish a baseline for future development in integrative genetic research.

## 2 Analytical strategies in colocalization analysis

We first consider two analytical strategies that differ in various implementations of SNP-level colocalization approaches. The first strategy concerns specification of enrichment levels of eQTLs in causal GWAS hits (i.e., the prior), and the second strategy is related to the modeling considerations for allelic heterogeneity that is wide-spread in molecular QTL data (i.e., the likelihood). We start by establishing the statistical connections and equivalence among different probabilistic colocalization methods.

### 2.1 Connections of existing probabilistic colocalization approaches

For a given genetic variant, the probabilistic quantification of colocalization is essentially to evaluate the conditional probability,

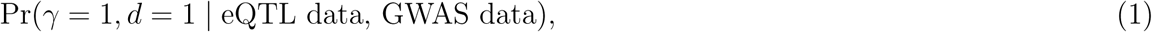

where indicators *γ* and *d* denote the latent causal association status of the target SNP with respect to the complex and gene expression traits of interest, respectively. All known probabilistic colocalization approaches aim to compute (1), which is carried out by applying the Bayes rule, i.e.,

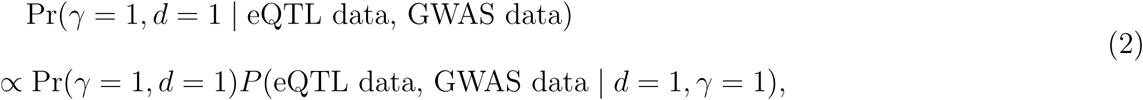

Noticeably, the computation would require an explicit specification of the prior probability Pr(*γ* = 1, *d* = 1).

The prior quantity, Pr(*γ* = 1, *d* = 1), reflects the frequency of colocalization sites in all interrogated variants and can be equivalently specified by the product of *p*_*d*_ := Pr(*d* = 1) and Pr(*γ* = 1 | *d* = 1). In ENLOC/fastENLOC, a set of enrichment parameters (*α*_0_, *α*_1_) are introduced to parameterize Pr(*γ* = 1 | *d*), i.e.,

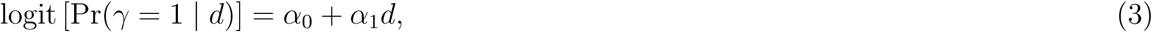

where *α*_1_ is the log odds ratio quantifying the enrichment level of molecular QTLs in GWAS hits. Note that the marginal association probabilities *p*_*d*_ and *p*_*γ*_ := Pr(*γ* = 1), representing the frequencies of causal eQTL and GWAS hits, are typically easy to estimate from the data [15, 16] and have general consensus.

In the implementation of *coloc*, the priors are defined by *p*_1_ := Pr(*γ* = 0, *d* = 1), *p*_2_ := Pr(*γ* = 1, *d* = 0), and *p*_12_ := Pr(*γ* = 1, *d* = 1). The equivalent parametrization by (*p*_*d*_, *α*_0_, *α*_1_) is given by

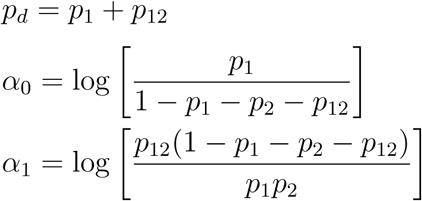

The third approach, eCAVIAR, makes a simplifying assumption,

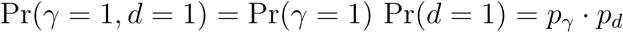

Note that this prior independence assumption is rather strong. The equivalent presentation in ENLOC parameterization is given by

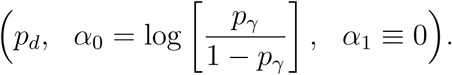

Given the equivalence of different formulations in all methods, our subsequent discussion will focus on the ENLOC/fastENLOC parameterization because of its convenient interpretation. It is important to note that, when *p*_*γ*_ and *p*_*d*_ are estimated from the observed data, the combination of (*α*_0_, *α*_1_) becomes constrained (Section 1 of the Supplementary Material). Specifically, the enrichment parameter, *α*_1_, becomes the only free parameter and needs explicit estimation/specification.

### 2.2 Specification of enrichment parameter

Provided that the prior specification is inevitable, we first perform a series of analyses to evaluate the sensitivity of the outcomes from colocalization analysis to the enrichment parameter. To isolate the effect of the enrichment parameter *α*_1_, we only consider computing the colocalization probability for a single SNP. Particularly, we consider weak, modest, and strong association evidence from respective eQTL or GWAS analysis, and examine the effect of varying enrichment levels on the magnitude of resulting SNP-level colocalization probabilities. The details of the analytical derivation are shown in the Methods Section and Section 2 of the Supplementary Material, and the range of the enrichment values is informed by our real data analysis presented in Section 4.

The results are summarized in Figure 1. In general, SNP-level colocalization probabilities are sensitive to the enrichment prior specification. However, depending on the combination of strength of evidence from individual association studies, different combination categories are differentially impacted. Specifically, when the eQTL and GWAS association evidence are both strong or both weak, the resulting colocalization probabilities are relatively stable with respect to the changes of the enrichment prior (in a wide and meaningful range). This phenomenon can be intuitively explained: when the marginal association evidence is highly informative (including the case that association evidence is weak, i.e., the evidence for no association is strong), the likelihood for colocalization is overwhelming and the prior impact is diminished. On the other hand, SNPs with modest association evidence from either GWAS or eQTL analysis seem most sensitive to the prior specification, as a strong enrichment assumption can significantly increase the corresponding probability of colocalization.

**Figure 1:**
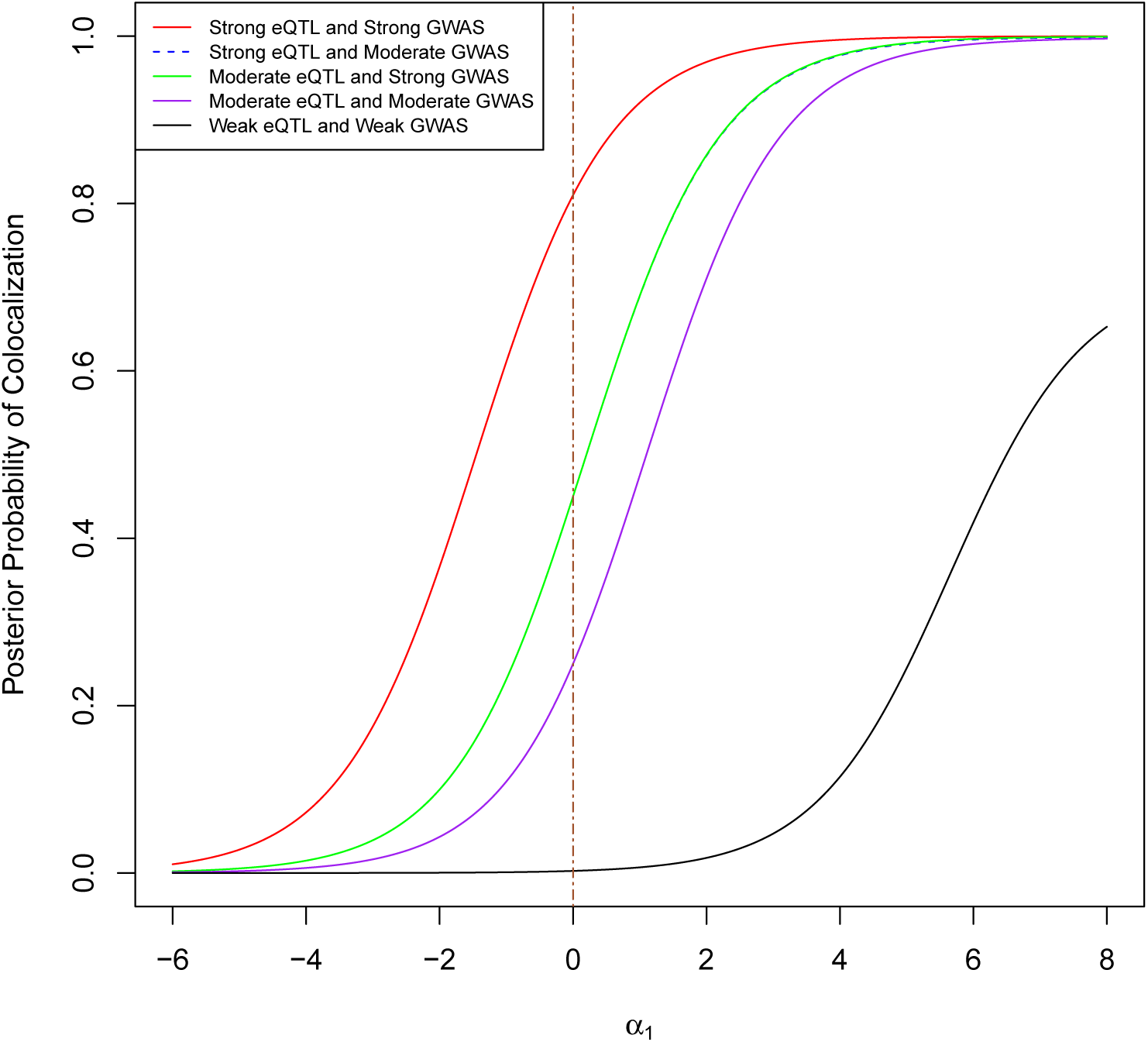
The impact of pre-defined enrichment level (*α*_1_) on the SNP-level colocalization probability. The different curves represent different combined levels of marginal association evidence for a particular SNP from the eQTL and GWAS studies. The range of the *α*_1_ values is informed by our real data analysis of 4,091 GWAS traits and the eQTL data from the GTEx project.

From this numerical experiment, we conclude that all colocalization methods likely agree on colocalization sites that show strong association evidence in both traits. However, the sensitive nature of the outcome with respect to the prior specification should caution practitioners. Because colocalization analysis is often treated as a discovery process similar to a hypothesis testing procedure, false positives, i.e., type I errors, should be carefully guarded against. From the observation of the numerical experiment, aggressively setting a high enrichment value tends to flag many more colocalization sites than setting a conservative value. However, this is also dangerous for inflating the type I error rates. Thus, we conclude that a conservative enrichment prior is, in principle, acceptable and may be preferable. On the other extreme, simply setting *α*_1_ = 0 in all circumstances can be too conservative and lead to severe loss of power.

For an ideal Bayesian analysis, the prior information should be derived from historical analyses of similar types. In the case that such historical information is unavailable, ***we recommend to estimate required hyper-parameters from the observed data***. Recently, [17] proposes to perform sensitivity analysis of the priors for identified colocalization signals. While we completely agree that understanding prior sensitivity is critical for practitioners, it should be noted that sensitivity analysis alone does not provide a justification for selecting a specific set of priors. Because the true association status for either trait is not observed, the estimation can be intrinsically challenging. The multiple imputation procedure implemented in ENLOC/fastENLOC, designed specifically for dealing with the “missing” association status, has shown the ability to provide robust and reliable enrichment estimates.

### 2.3 Accounting for allelic heterogeneity

Allelic heterogeneity (AH) refers to the phenomenon of distinct genetic variants at a locus simultaneously affecting the same phenotype. The analytical consideration for AH relates to the likelihood computation in (2) and can have a significant impact on the outcomes of colocalization analysis. In the analysis of eQTL and GWAS data, a locus is typically defined as the cis region of a target gene, e.g., a 2Mb window centered around the transcription start site [18]. At the scale of such loci, AH is a widespread phenomenon in gene regulations based on the strong evidence from recent eQTL data [19, 18]. Others have also reported that multiple independent causal GWAS hits can co-exist within such genomic regions [20]. Some colocalization methods, e.g., coloc, JLIM, are built upon the assumption that there is at most one causal SNP within a locus for a given trait, i.e., the “one causal variant” (OCV) assumption. This assumption primarily provides computational convenience in computing the likelihood of colocalization: if the OCV holds, the LD information between candidate SNPs within the locus becomes obsolete for the computation [21, 15, 22]. Computational methods based on this assumption have been shown successful in many different types of association analysis for a single trait, but the assumption has not been systematically investigated in the context of colocalization analysis. Given the widespread AH that is commonly observed in recent molecular QTL data, the OCV is likely often violated. Thus, we are interested in exploring its implications on both false-positive and false-negative errors in colocalization analysis.

We first conduct simulation studies using real genetic data from the GTEx project and simulate gene expression and complex trait data based on linear regression models. To isolate the effect of AH, we focus on simulating strong genetic effects for all eQTLs and GWAS hits. (As shown in the previous section, these signals are robust to the misspecification of enrichment parameters.) Furthermore, both expression and complex trait data are independently generated from the same genotype data, which ensures LD mismatch is not a factor in the analysis. Our simulation considers the following scenarios for each gene-trait pair within a genomic locus:

1. single causal variants in both eQTL and GWAS data, no colocalization
2. AH in eQTL data, a single causal variant in GWAS data, no colocalization
3. single causal variants in both eQTL and GWAS data, single colocalization event
4. AH in eQTL data, a single causal variant in GWAS data, single colocalization event

Note that the first two scenarios are designed to investigate potential false positive findings and the last two are for false negative findings. The assembled data for analysis consist of different mixtures of the four scenarios, such that we could evaluate both the type I and the type II errors. The simulation details are provided in the Method section.

We use two different methods to analyze the simulated data. Particularly, we apply the default *coloc* method to represent methods making the OCV assumption and without explicit modeling of AH. We apply fastENLOC to represent methods that explicitly model potential AH. The results of our simulations are in Table 1. In all of our simulated scenarios, we find that both types of approaches control the type I errors. However, when AH is presented, the power of *coloc* is approximately half of the power of fastENLOC. The relative ratio of power between the two types of approaches is expected. When two independent eQTL signals co-exist, *coloc* identifies one signal with a stronger signal-to-noise ratio as the sole casual eQTL, which leads to a false-negative finding when the unselected eQTL overlaps the causal GWAS hit.

**Table 1:**
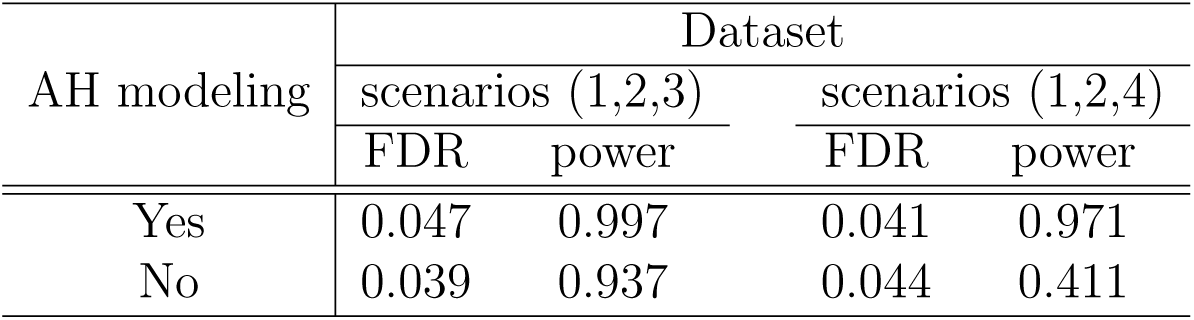
The impact of modeling consideration of AH on FDR and power in colocalization analysis.

| AH modeling | Dataset |  |  |  |
| --- | --- | --- | --- | --- |
|  | scenarios (1,2,3) |  | scenarios (1,2,4) |  |
|  | FDR | power | FDR | power |
| Yes | 0.047 | 0.997 | 0.041 | 0.971 |
| No | 0.039 | 0.937 | 0.044 | 0.411 |

Even though the simulation study does not indicate apparent inflation of type I errors, theoretical analysis shows that there are potential false positives directly related to the simplifying assumption of no AH. There are at least two concerns in making the OCV assumption in colocalization analysis. First, some implementations of the assumption, e.g., *coloc*, enumerate all possible causal association configurations from both traits to compute the normalizing constants and desired colocalization probabilities. Under the simplifying assumption of OCV, there are (*p*+1)^2^ possibilities precisely (where *p* represents the number of SNPs in the locus). If the assumption is violated, many more necessary scenarios are uncounted (the total possibilities without constraints are 2^*p*^). This factor can lead to under-estimation of the normalizing constants and over-estimation of the colocalization probabilities (Supplementary Figure S1). Second, false positives can be carried over from single-SNP analysis in the marginal studies. It is known that some non-causal SNPs that are in partial LD with multiple causal SNPs can generate the most significant single-SNP association evidence [19]. Such false-positive findings based on single-SNP association evidence are maintained and carried over into the colocalization analysis under the OCV assumption. Consequently, it becomes a source of false-positive colocalization findings. In contrast, methods that explicitly perform multi-SNP fine-mapping analysis can effectively dissect different scenarios by accounting for LD. Hence, they are unlikely to suffer from such false-positive findings. In summary, we conclude that the OCV assumption can potentially lead to anti-conservative quantifications of colocalization probabilities. Although the extent of the anti-conservativeness may not lead to observable inflations of type I errors within our simulations, its effect can be observed in the numerical experiments examining the calibration of the reported colocalization probabilities (Section 3 of the Supplementary Material).

More generally, ***our recommendation for dealing with likelihood computation in colocalization analysis is to apply dedicated fine-mapping approaches, and separate fine-mapping and colocalization analysis.*** Such a strategy has been established in eCAVIAR and fastENLOC. We note that state-of-the-art of fine-mapping approaches [16, 9, 23] all have the ability to account for AH without making the OCV assumption. It is also intuitive that the colocalization analysis should be based on the best possible fine-mapping results. This is because the inaccuracy from the fine-mapping analysis will inevitably translate into inaccuracy in subsequent colocalization.

## 3 Practical factors in colocalization analysis

There are many practical factors in colocalization analysis that analysts have little control over. Nevertheless, their impacts on the outcomes are profound. Having established the two fundamental inference principles, we proceed to assess the empirical performance of colocalization analysis and investigate the other performance-impacting factors using realistically simulated eQTL and GWAS data.

### 3.1 Empirical assessment of probabilistic colocalization analysis

To construct a simulated dataset that resembles real applications of colocalization analysis, we simulate 20,000 non-overlapping genes with 1500 SNPs within each cis-region. The scale of the simulated datasets resembles real applications of genome-wide colocalization analysis. We use the real genetic data from 400 participants of the GTEx project. In order to circumvent the issue of LD mismatch in this particular experiment, we use the same set of genotype data to simulate both the expression and complex trait phenotypes. Within each *cis* region, the causal eQTLs are randomly selected from a series of independent Bernoulli trials, such that on average there are three causal eQTLs per gene (i.e., *p*_*d*_ = 2 *×* 10^−3^). Similarly, we sample causal GWAS SNPs conditional on the simulated eQTL status using the probability model (1) with a true *α*_1_ value of 4. As a result, the simulated data set consists of 2,103 colocalized association signals distributed in 2,001 unique genes. Given the truly causal SNPs for each gene, we independently generate the molecular and complex trait data for the 400 individuals based on standard multiple linear regression models. Specifically, the genetic effects of each causal variant-trait pair is independently drawn from the distribution *N* (0, 1). The residual errors for each trait and each individual is also independently generated from the standard normal distribution.

To analyze the simulated dataset, we first perform separate Bayesian fine-mapping analysis for the simulated eQTL and GWAS datasets using the software package DAP-G [16]. Utilizing the resulting probabilistic annotations, we apply fastENLOC to estimate the enrichment parameters with the default shrinkage setting. As expected, the estimated enrichment parameter *α*_1_ is slightly under-estimated, however reasonably close to the true value (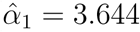 with the standard error 0.039). We then perform colocalization analysis using fastENLOC with the estimated and true enrichment parameters, respectively. For comparison, we also run *coloc* with its default prior model parameters and the true parameters, respectively. Note that the differences between *coloc* and fastENLOC results based on the true enrichment parameters should reflect the difference in fine-mapping analysis, including the consideration of AH.

We first examine the false positive rate and the power at 5% FDR level for different analysis settings (Table 2). We find that severe inflation of type I errors only occurs at the *coloc* run with its default model priors (which significantly exceeds the true enrichment parameter). All other analysis settings, including the *coloc* run with the true enrichment parameters, show proper control of the desired false discovery rate. For the methods that control the type I errors, the power seems low across the board (Table 2). The under-estimation of the enrichment parameter, *α*_1_ due to shrinkage only explains a small fraction of the power loss. Although the power and type I error analysis only focus on high values in colocalization quantifications, our conclusion extends to the full probability spectrum: additional inspection on the calibration of the regional colocalization probabilities (RCPs) also confirms that various methods with justified or true priors yield conservative colocalization results (Supplementary Figure S1).

**Table 2:** Realized false discovery rates and power of colcoalization analysis in simulated data. The * symbol indicates that the realized FDR is severely inflated from the control level (5%). The corresponding power is annotated by †. The table provides an overall assessment of various colocalization approaches. We observe that the type I errors are typically under control when the enrichment priors are justified. The power (for methods properly controlling FDR) is overall quite low in this setting that resembles realistic applications.

| Method | Number of discoveries | Realized FDR | Power |
| --- | --- | --- | --- |
| fastENLOC (estimated prior) | 406 | 0.034 | 0.186 |
| fastENLOC (true prior) | 458 | 0.046 | 0.208 |
| coloc (default prior) | 472 | 0.258* | 0.175<sup>†</sup> |
| coloc (true prior) | 200 | 0.045 | 0.095 |

To explain the loss of power, we identify two primary sources of false negative errors by an indepth examination of the simulated data and the corresponding analysis results. We refer to these two sources as class I and class II false negative errors in colocalization analysis. Specifically, we define

1. **Class I false negatives**: lack of power in association analysis of individual traits
2. **Class II false negatives**: inaccurate quantification of association evidence at SNP level for individual traits

The class I false negatives (FNs) represent the cases of failure in detecting at least one type of association signals (eQTL or GWAS) in genetic association analysis. The class II FNs represent the scenarios where both types of association signals are correctly uncovered at locus level, but the inaccurate SNP-level quantifications imply that the causal variants for the two types of traits are unlikely overlapping. In our simulation studies, 59.0% (1240) and 22.0% (463) of the true colocalization signals fall into the class I and II false negatives categories, respectively. To better visualize the two classes of FNs, we plot the true eQTL and GWAS effects of the colocalized signals with their corresponding labelled categories based on the fastENLOC results in Figure 2. Most points representing class I FNs (grey) are closely located around the axes, indicating at least one of genetic effects (eQTL or GWAS) is too small to be detected by the corresponding association analysis. Marginally at the 5% FDR level, the power for GWAS and eQTL is 44% and 62%, respectively. In comparison, most points representing class II FNs (cyan) and detected signals (red) are scattered around the two diagonals.

**Figure 2:**
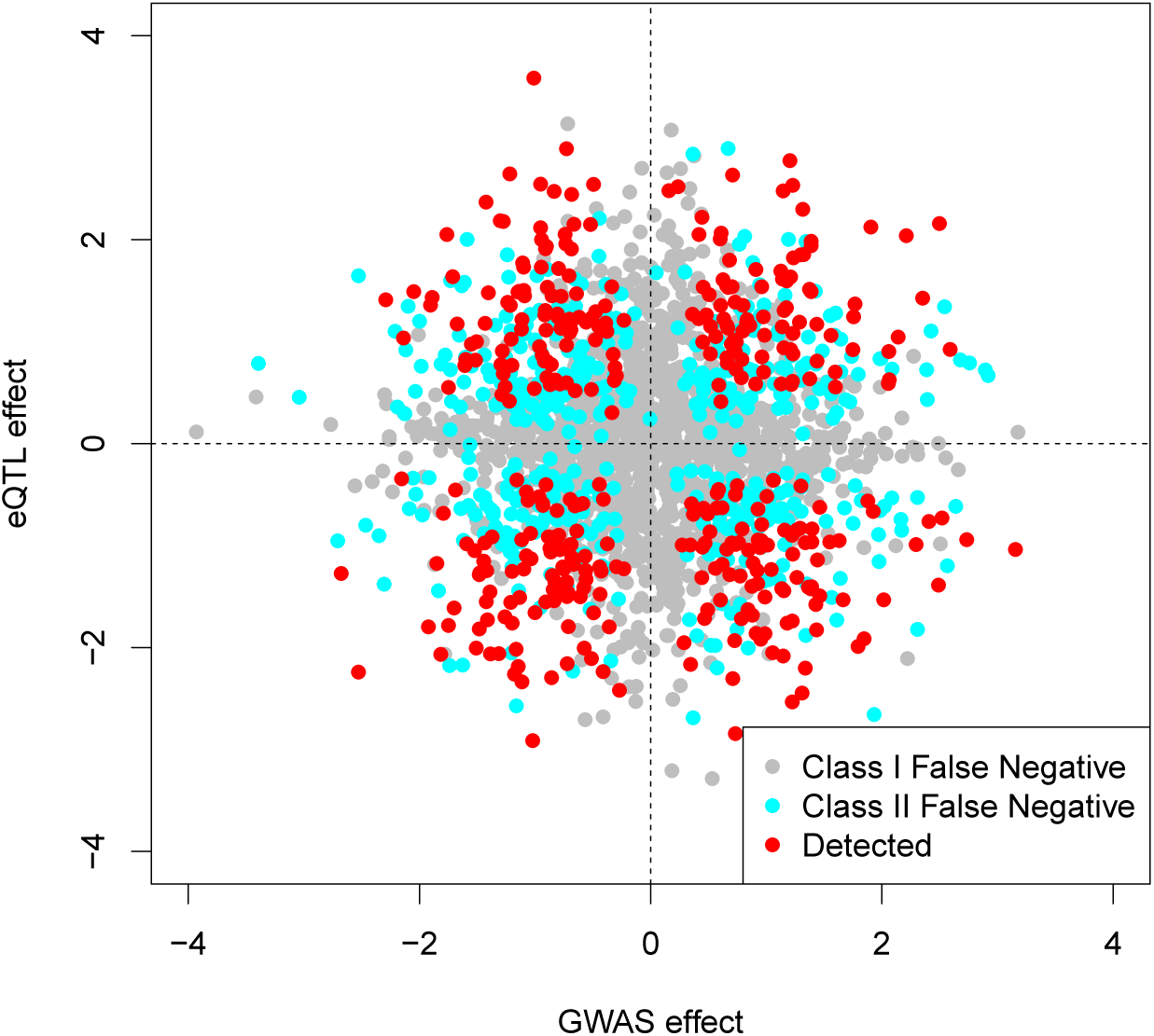
Classification of all Colocalized SNPs. All truly colocalized variants from our realistic simulations, classified as either a class I false negative, class II false negative, or successfully detected by fastENLOC. We see the expected pattern that most points near one of the axes are class I false negatives, while points far away from both tend to be detected by fastENLOC.

The impacts of class I FNs are easy to understand and well expected. As neither eQTL mapping nor genetic association analysis of complex traits achieves high power in practice, this class of FNs remains a primary source for failures in identifying colocalization sites.

A proportion of the class II false negatives can be explained by a “threshold” effect. For example, some modest eQTL signals barely clear the bar to qualify for significant eQTL findings, but the underlying evidence is not strong enough to ensure a significant colocalization discovery. Additionally, the class II FNs can occur even when the association signals for GWAS and eQTL can both be narrowed down into the same genomic locus with high confidence. This is because, at the SNP-level, it remains difficult to pinpoint the causal variants for both traits due to the combination of LD and insufficient sample size. The phenomenon of class II FNs are also closely related to the well-known fact in fine-mapping analysis: the lead (i.e., the most significant) SNPs that emerge from association analysis may not be the true causal SNPs [24, 25]. Incidental correlation between the genotypes of non-causal SNPs (in LD with the true causal variant) and residual errors from the outcome variable could lead to stronger empirical correlation, especially with limited samples. There is generally a higher level of mismatching between lead and causal SNPs when the underlying studies are underpowered. In any association analysis, Bayesian or frequentist, the lead SNPs are always regarded as the most plausible causal SNPs from the data. When the mismatch of lead and causal SNPs occurs in at least one trait, all algorithms are led to believe there is a lack of evidence for *SNP-level* colocalization, even though the signal clusters for both traits are correctly identified.

To provide a visualization, we compute a PIP ratio for the causal SNP versus the lead SNP (causal-vs-lead PIP ratio) in each signal cluster harboring a true colocalized signal for both simulated eQTL and GWAS data. The ratio = 1 indicates that the lead SNP is indeed the causal SNP (or they are in perfect LD). We further compute a combined ratio by multiplying the two trait-specific causal-vs-lead PIP ratios for each signal cluster. Note that the combined ratio = 1 suggests the causal SNPs are identified as lead SNPs in both traits, whereas the combined ratio < 1 indicates that in at least one trait, the lead SNP and the causal SNP do not match. The comparison of various PIP ratios between detected and class II false negative signals are shown in the histograms of Figure 3. The overall patterns in Figure 3 indicates that in detected colocalization signals, the vast majority of causal SNPs are indeed lead SNPs in both traits; many mismatches between causal and lead SNPs lead to false negatives in identifying the colocalized signals.

**Figure 3:**
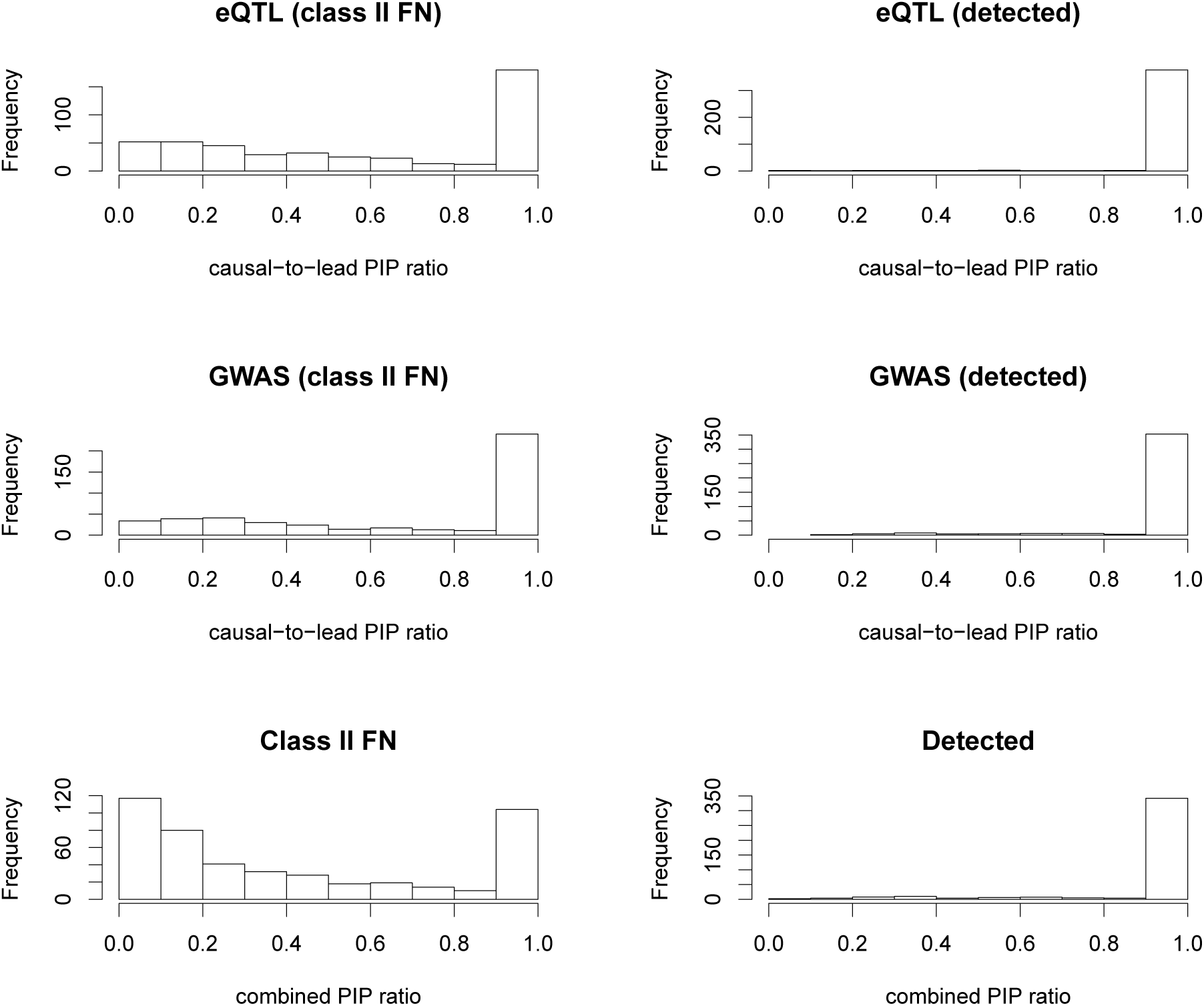
PIP ratios for all signal clusters. The first column of histograms represent the ratio of causal SNP PIP to lead SNP PIP for all class II false negatives from the eQTL simulated dataset, the GWAS dataset, then both combined. The second column represents the same ratio, but among all successfully detected colocalizations from the respective datasets.

Both classes of false negative errors are intrinsic to genetic association analysis and well-known. However, it is somewhat surprising to observe that the combined effects from these factors have such a drastic effect on the power of colocalization analysis – even when the association analysis for individual traits is considered well-powered.

In practice when truth is unknown, the analyst can still assess the relative power of the colocalization analysis based on the estimates of *p*_*d*_, *p*_*γ*_, and *α*_1_. The expected number of colocalization sites based on the enrichment analysis can be computed by 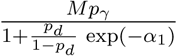, where *M* represents the total number of genetic variants. In this simulation, the fastENLOC estimates

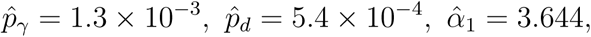

and the expected number of the colocalization sites is ∼ 800, which represents a lower bound estimate of true colocalization sites (due to the conservative estimate of *α*_1_ and the class I FNs). The number of detected sites at 5% FDR level is roughly half of the expected sites. Henceforth, we refer to this proportion as the rejection-to-expectation (rej-to-exp) ratio, which represents an upper-bound estimate of the empirical power.

### 3.2 Mismatching LD structures

Most existing colocalization analysis approaches are built on the experimental scheme known as the *two-sample* design, where the eQTL and GWAS data do not share common samples. While this design allows for using valuable eQTL resources, e.g., GTEx data, to analyze a wide range of GWAS data collected from many different cohorts, it raises some practical concerns. To our knowledge, all existing methods implicitly assume that the LD structures between the two association samples are identical, which is at best questionable when the two sets of association data are collected from different cohorts. In general, there is a lack of empirical evaluation on how different levels of mismatching between LD structures affect the outcomes of colocalization analysis. To address this issue, we design a simulation experiment utilizing multi-population genetic data to quantify the effects of mismatching LD patterns on colocalization analysis with two-sample designs.

We take the genetic data from the GEUVADIS project, which consists of samples from four European populations: CEPH (CEU), Toscani (TSI), British (GBR) and Finns (FIN); and one African population, Yoruban (YRI) [26]. We select the SNPs located within a 200kb *cis*-region from 6,977 protein coding and lincRNA genes, each of which contains at least 500 candidate SNPs. For each gene, we first sample the causal eQTL and GWAS SNPs from its candidate cis-SNPs. We then simulate a single eQTL data set using the FIN data. We subsequently generate 5 GWAS datasets using the genotype data and pre-determined GWAS association status for all 5 population groups. Note that for the Finnish population, the LD patterns are perfectly matched for GWAS and eQTL analysis, which forms a baseline for evaluating the effects of LD mismatching.

We analyze the 5 pairs of eQTL-GWAS data using fastENLOC. Our comparisons focus on the enrichment estimates, false positive colocalization findings and the power. The results are summarized in Table 3. In all cases, we do not observe any inflation of false positive colocalization findings – the type I error rates are properly controlled in all datasets. The impact of LD mismatch is reflected by the under-estimation of the enrichment parameters and the diminished power, especially in noting that the power of GWAS discovery (which is perfectly correlated with estimated *p*_*γ*_) are overall close in all populations. In the extreme case of mismatch, i.e., the analysis of YRI GWAS data and FIN eQTL data, we find that the enrichment parameter *α*_1_ is most severely underestimated, and the resulting power is reduced to 50% of the perfectly matching association data (i.e., FIN GWAS and FIN eQTL). We also note that the underestimation of the enrichment parameter only explains a small proportion of the loss: even when the true enrichment parameter is used, the power of colocalization from analyzing the YRI GWAS data remains significantly lower than the other European data sets. Within the European populations, the effects of LD mismatch on colocalization analysis are also noticeable. The comparison within the European populations may also be complicated by differences in sample sizes, where TSI and CEU have the largest (*n* = 92) and smallest (*n* = 79) sample sizes, respectively. The sample size difference is directly linked to the power of GWAS discovery.

**Table 3:**
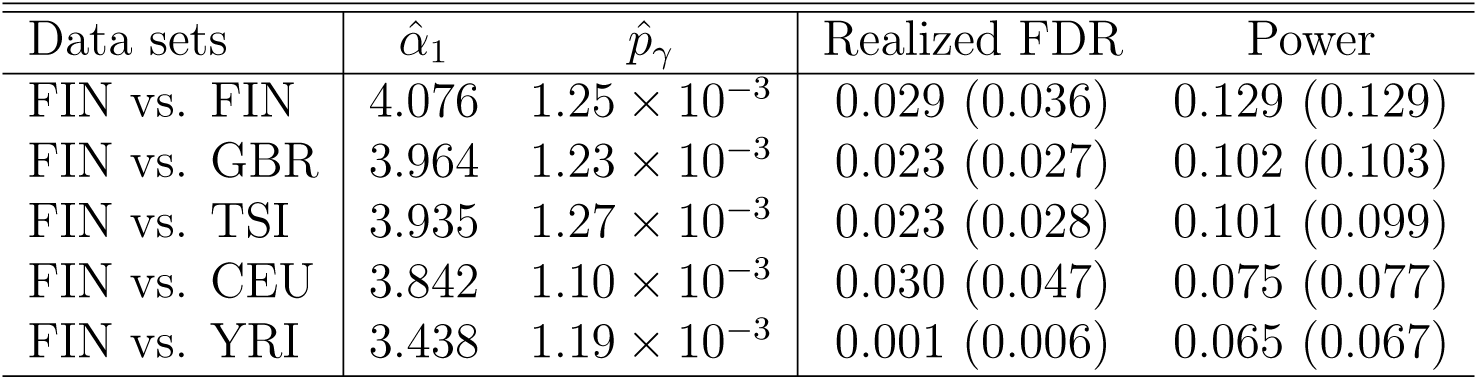
LD mismatch impact on enrichment estimation, FDR, and power. The enrichment estimates 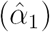 and from all combinations of eQTL and GWAS datasets we used for this *×* analysis. The estimated frequency of GWAS hits 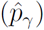 reflects the GWAS power of each GWAS dataset. (Note that true *p*_*γ*_ = 1.92 × 10^−3^.) The quantities in parentheses show the realized FDR and power when using the true enrichment parameter in the colocalization analysis.

| Data sets | $\hat{\alpha}_1$ | $\hat{p}_\gamma$ | Realized FDR | Power |
| --- | --- | --- | --- | --- |
| FIN vs. FIN | 4.076 | $1.25 \times 10^{-3}$ | 0.029 (0.036) | 0.129 (0.129) |
| FIN vs. GBR | 3.964 | $1.23 \times 10^{-3}$ | 0.023 (0.027) | 0.102 (0.103) |
| FIN vs. TSI | 3.935 | $1.27 \times 10^{-3}$ | 0.023 (0.028) | 0.101 (0.099) |
| FIN vs. CEU | 3.842 | $1.10 \times 10^{-3}$ | 0.030 (0.047) | 0.075 (0.077) |
| FIN vs. YRI | 3.438 | $1.19 \times 10^{-3}$ | 0.001 (0.006) | 0.065 (0.067) |

Overall, in relative terms, our observation suggests that the power loss suffered from the LD mismatching is qualitatively less severe than from the imperfect power of individual association analysis – as long as the eQTL and the GWAS samples are from reasonably close populations. In addition, the power loss caused by LD mismatching may be compensated by increased power in single-trait association analysis.

## 4 Colocalization analysis of 4,091 GWAS datasets and GTEx eQTL data

To provide a comprehensive summary of colocalization analysis using the current available GWAS and eQTL data, we analyze 4,091 complex trait datasets and the final release of the GTEx data (v8) from 49 tissues [18, 6]. In total, we perform colocalization analysis on 200,459 trait-tissue pairs using fastENLOC. The biological implications from the colocalization analysis, coupled with PrediXcan analysis [4], have been reported and discussed in [6]. In this section, we focus on the technical perspective of the colocalization analysis and provide a high-level summary of colocalization analysis over a wide range of complex traits with currently available GWAS and eQTL data sets.

We first examine the empirical distribution of the enrichment estimates,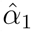, over the 200,459 trait-tissue pairs. The histogram in Figure 5 shows the empirical distribution of the enrichment estimates. We observe a clear bi-modal distribution: the estimates from the vast majority of the trait-tissue pairs are close to 0, and there is also a noticeable peak centered around *α*_1_ = 4.

**Figure 4:**
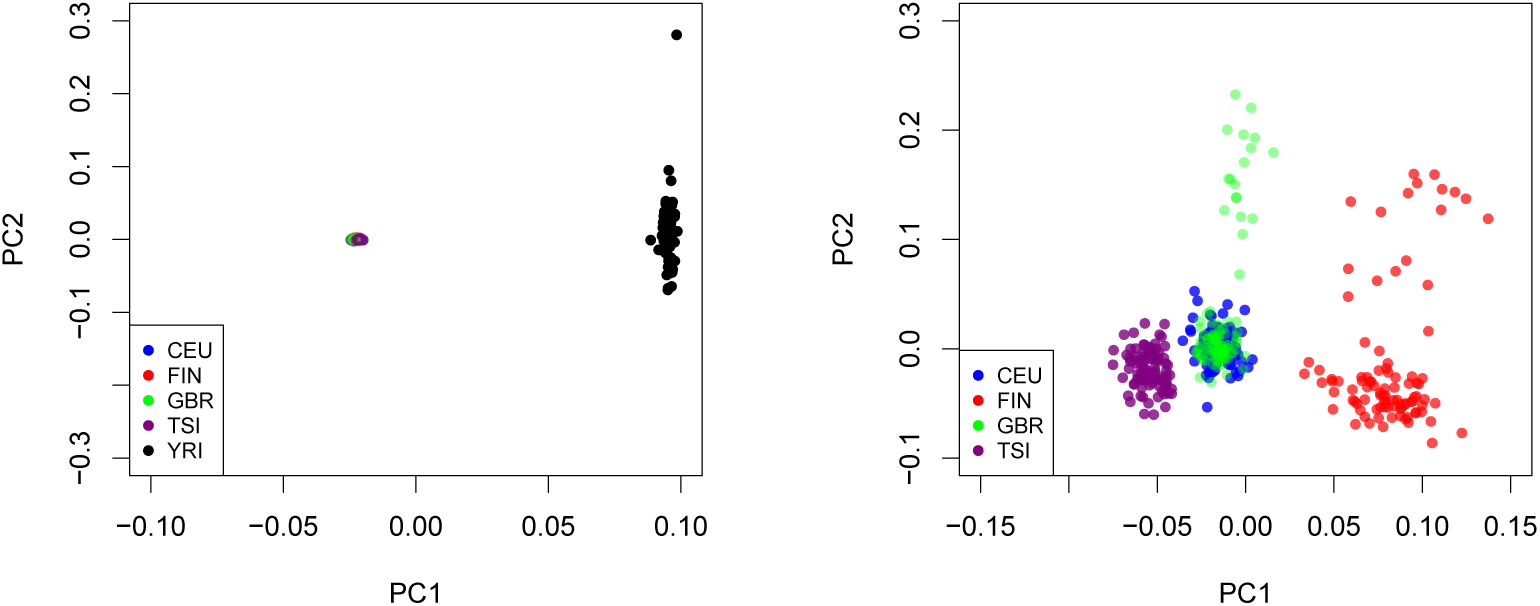
Population structures represented by PCA plots in GEUVADIS data. The left panel show the PCA plots (PC1 vs. PC2) with the samples from all populations. The right panel shows the PCA plots using European samples only. Based on these plots, we expect maximum LD mismatch between YRI and any European population.

**Figure 5:**
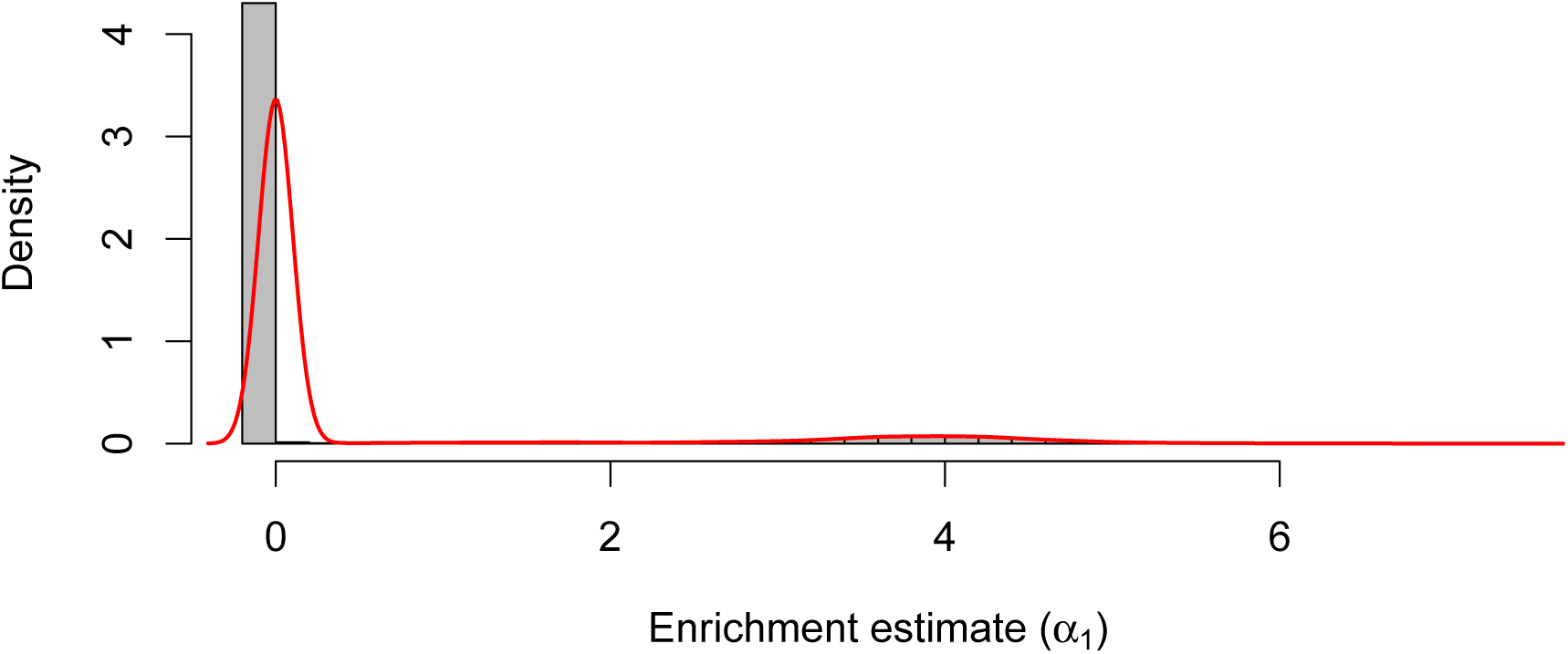
Enrichment estimates from all tissue-trait pairs in the phenomexcan analysis. The histogram displays a bimodal distribution, with a sharp peak at *α*_1_ = 0 and a wide peak centered around *α*_1_ = 4.

Next, we inspect the significant findings from the analysis of each individual trait-tissue pair. Based on the results from the enrichment analysis, we compute the expected number of colocalization sites for each trait-tissue pair. Additionally, we identify high-confidence colocalization sites at the 5% FDR level based on the output of RCP values using the Bayesian FDR control procedure. In total, 15,975 sites pass this type I error control threshold in all trait-tissue pairs. We consider the trait-tissue pairs with more than 50 expected colocalization sites as “well-powered”. For this set of trait-tissue pairs, we compare the expected colocalization sites and the identified high-confidence sites at 5% FDR level (Figure 6). The average rej-to-exp ratio in this set is 10.7% (median = 9.43%). Recall that rej-to-exp ratios represent the upper-bound estimates of empirical power for colocalization analysis. This result indicates that the current colocalization analysis is (severely) under-powered for most of the trait-tissue pairs.

**Figure 6:**
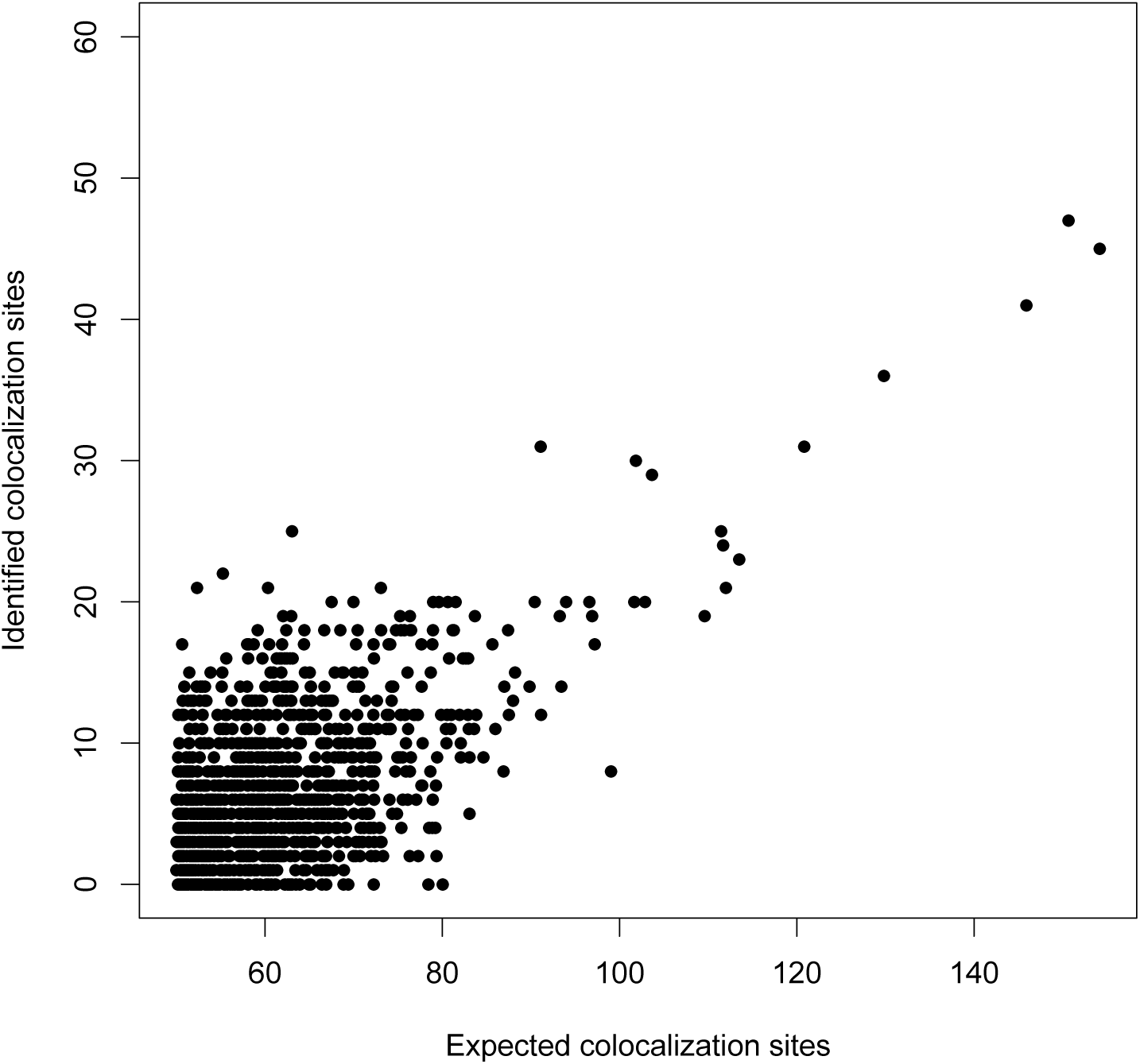
The comparison of the calculated expected colocalization sites and the detected high-confidence sites, among well-powered trait-tissue pairs. This apparent lack of power falls in line with our findings from our simulations.

## 5 Discussion

In this paper, we have systematically explored both the analytical and the practical factors that impact the performance of probabilistic colocalization analysis for a molecular and a complex trait. We identify a single analytical factor, i.e., the specification of prior enrichment levels, that can lead to significant inflation of false-positive findings, and we recommend to estimate the critical enrichment parameters directly from the data. On the other hand, we find that a combination of analytical and practical factors, including modeling considerations for AH, LD mismatch, and imperfect power in association analyses, could severely diminish the power of SNP-level colocalization discoveries. As a result, current approaches often fail to identify the majority of colocalization signals in practical applications, even when they are appropriately applied. We argue that understanding the promise and limitations of the current state-of-the-art is critical for the practitioners to properly anticipate and correctly report their findings. For colocalization analysis of currently available molecular QTL and GWAS data, we may need to embrace the noticeable discrepancy between “expected colocalized signals” and the actual identified “significant colocalization findings”.

There are many ways in which we can improve the power of existing colocalization methods based on our findings in this paper. For example, analytical strategies in improving the enrichment estimation, applying better fine-mapping methods, and explicitly modeling varying LD patterns across datasets will most likely result in enhanced power. Nevertheless, we suspect that improving the quality of the GWAS and molecular QTL datasets should have a more direct and visible impact. We note that most existing molecular QTL studies are limited by the high-throughput phenotyping cost and have modest sample sizes and relatively high-levels of experimental noise. The state-of-the-art eQTL annotations generated by the GTEx project are derived from bulk tissues of < 1, 000 samples. The current technology advancement, e.g., applying single-cell technology for molecular QTL mapping, combined with proven statistical strategies for data aggregation, e.g., a meta-analysis of molecular QTLs, could significantly enhance colocalization discoveries.

There are other analytical issues in colocalization analysis that we do not discuss in the main text. For example, some practitioners perform LD pruning on a candidate region before a formal colocalization analysis. We do not recommend such a practice. This is simply because the pruning process has the effect of “collapsing” linked genetic variants into a single proxy variant. For SNP- level colocalization analysis, this could potentially introduce false-positive colocalizations. As we have shown in the paper, SNP-level colocalization analysis is an application of Bayesian inference. Thus, the analytic strategies should follow the principles of Bayesian inference. Particularly, practitioners should vigorously justify the prior specification in context-specific applications. “Borrowing” priors from existing literature should be cautioned unless the circumstances are extremely similar. More generally, we recommend estimating the required hyper-parameters directly from the observed data. For likelihood computation, we strongly recommend separating the analysis of fine-mapping and colocalization. This strategy enables the best computational methods to generate accurate probabilistic annotations for GWAS hits and molecular QTLs, which should improve the power of subsequent colocalization analysis. We acknowledge that our recommended analytical strategies tend to sacrifice some computational convenience, but the gain in power and accuracy warrants the efforts.

Although our discussions in this paper are exclusively illustrated using two complex traits, the general principles extend to the analysis of multiple traits [27, 28]. All the analytical and practical factors that we have discussed take effects in the SNP-level colocalization analysis for more than two traits. If not adequately dealt with, the resulting adverse effects can be even more severe. For example, both classes of false-negative errors discussed in Section 3.1 increase with more traits considered. Additionally, the existing enrichment estimation procedure via multiple imputation does not scale well regarding a large number of traits (i.e., *≥* 5). Therefore, extending the best practice of colocalization analysis from two traits to multiple traits remains a critical challenge.

Colocalization analysis is also connected to other types of integrative analysis approaches, e.g., transcriptome-wide association studies (TWAS). In analyzing eQTL and GWAS data, TWAS utilizes the same sources of input data as the colocalization analysis. Its results, however, have some unique causal implications provided that a set of assumptions is met [29]. Despite the difference in their theoretical origins, positive findings from the two analyses can be driven by similar types of signals. Recent studies [6] find that integrating colocalization analysis into TWAS can improve its sensitivity and specificity. More generally, the two prevailing types of integrative analysis approaches can complement each other. Thus, further exploration of their connections and distinctions becomes an important future direction.

## 6 Methods

### 6.1 Overview of existing probabilistic colocalization approaches

#### coloc

*coloc* is first proposed in [12]. It makes the “one causal variant” (OCV) assumption for each trait in each candidate region. Under this specific assumption, it enumerates five possible distinct models/hypotheses within a region. These models are: 1) there are no causal variants from either trait (*H*_0_); 2) there is only a causal eQTL variant (*H*_1_) but no causal GWAS variant; 3) there is only a causal GWAS variant but no causal eQTL variant (*H*_2_); 4) there are distinct causal SNPs for both eQTL and GWAS, (*H*_3_); and 5) there is a colocalized signal (*H*_4_). For each hypothesis, the corresponding posterior probability is computed by considering all compatible latent association status configurations from GWAS and eQTL data via Bayesian model averaging (BMA). The colocalization within each region is quantified by the posterior probability of *H*_4_ (PPH4).

Under the OCV assumption, the likelihood functions for all possible association scenarios can be analytically computed from the single-SNP association statistics, e.g., *z*-statistics, without the need of explicit modeling of LD [15]. The implementation of coloc allows user-specified priors for SNP *i*, i.e.,

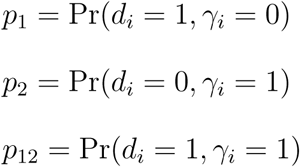

Recall that *d*_*i*_ = 1 indicates that SNP *i* is a causal eQTL and *γ*_*i*_ = 1 indicates that SNP *i* is a casual GWAS variant. By default, *p*_1_ and *p*_2_ are set to 10^−4^, and *p*_12_ is set to 10^−5^, which corresponds to

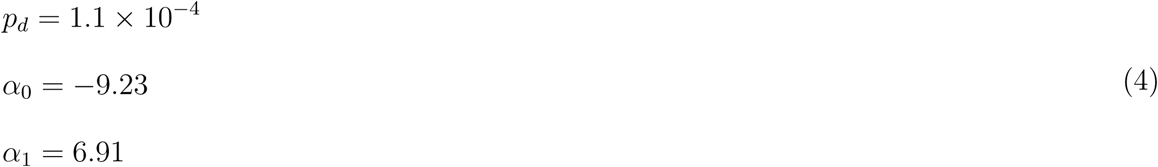

in the formulation of fastENLOC priors. From the observed data (Figure 5), *α*_1_ close to 7 represents a relatively high enrichment level.

It is worth pointing out that *coloc* requires only summary-level statistics from single-SNP association analysis as input and performs fine-mapping internally assuming OCV. This feature certainly brings desired computational convenience and efficiency.

#### eCAVIAR

eCAVIAR, proposed in [13], is built upon a sophisticated Bayesian multivariate fine-mapping algorithm, CAVIAR [23]. The CAVIAR algorithm enables computation of Pr(*d*_*i*_ = 1 | eQTL data) and Pr(*γ*_*i*_ = 1 | GWAS data) based only on summary-level statistics from single-SNP association analysis and an LD matrix between candidate SNPs. Based on the marginal fine-mapping result, eCAVIAR computes a SNP-level colocalization posterior probability, or CLPP, by

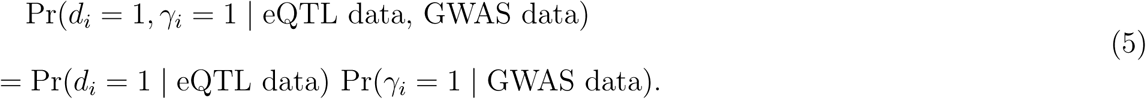

As we previously discussed, this equation corresponds to the special case that assumes *α*_1_ = 0, i.e., there is no enrichment of eQTL signals in causal GWAS hits. Based on the evidence observed from the real data (Figure 5), this assumption is conservative for a subset of complex traits.

Notably, eCAVIAR only provides a colocalization quantification at the SNP level but not for a genomic region. This is also a distinction between eCAVIAR and other approaches.

#### fastENLOC/ENLOC

ENLOC is first proposed in [14]. An improved version, fastENLOC, with accelerated computation and enhanced precision is recently described in [6]. A distinct feature of ENLOC/fastENLOC is its embedded function of estimating the enrichment level of eQTLs in GWAS hits, i.e., (*α*_0_, *α*_1_), directly from the data. By treating the latent vectors of causal association status, ***γ*** and ***d***, as missing data, ENLOC employs a multiple imputation strategy and an EM algorithm for enrichment estimation. Given the enrichment estimates, ENLOC applies an empirical Bayes framework to compute the SNP-level colocalization probability strictly based on the Bayes rule, in which Eqn. (5) of eCAVIAR becomes a special case.

fastENLOC also takes advantage of the concept of credible sets of independent association signals inferred from multi-variant fine-mapping analysis to compute a colocalization probability for each LD region. A Bayesian credible set consists of a group SNPs in LD that represent the same underlying association signal. By utilizing credible sets, fastENLOC ensures correct dependence and independence structures in the sampling procedure of the multiple imputation scheme for enrichment estimation. Additionally, the credible sets naturally form regional LD units for regional-level colocalization analysis. This is because even if we are unsure which exact variant represents the colocalized signal, we can evaluate the probability that *one* of the member SNPs in the credible set is causal for both GWAS and expression traits by Bayesian model averaging.

Similar to eCAVIAR, fastENLOC requires dedicated fine-mapping algorithm to generate required probability quantifications of marginal association evidence. Currently, DAP [16] and SuSiE [9] are two suitable methods for preparing fastENLOC input, because both have the ability to report signal clusters/credible sets in addition to traditional SNP-level posterior inclusion probabilities (PIPs). Additionally, both DAP and SuSiE work with summary-level statistics in a similar fashion as CAVIAR and eCAVIAR.

#### Other approaches

There are other available methods proposed for general colocalization analysis between multiple complex traits. The majority of methods in this category deal with a slightly different problem that does not require quantification of colocalization at the SNP-level. For example, both RTC (Regulatory Trait Concordance) [2] and JLIM (Joint likelihood mapping) [11] use formal hypothesis testing procedures to examine if a group of SNPs in LD contain both a causal eQTL and a causal GWAS hit. However, within the SNPs that are tightly linked, they are not able to distinguish if the two signals are overlapping at a single SNP. That is, there is no stringent distinction between the *H*_3_ and the *H*_4_ models in the formulation of *coloc*. Another approach, SMR [30], is built upon the framework of instrumental variable analysis and identifies SNPs that are associated with both GWAS and expression traits. The authors attempt to further distinguish colocalization (*H*_4_) from linkage (i.e., *H*_3_) by a hypothesis testing procedure named HEIDI [31]. Although the idea is intuitive, the HEIDI procedure is set up to claim colocalization by *accepting* the null hypothesis. As a result, rigorous quantification or proper control of false positive findings is lacking.

Finally, we want to emphasize that, despite the difference in their implementations, there is strong uniformity across all three probabilistic colocalization approaches: they share the same mathematical foundation and inference principles. Particularly, eCAVIAR can be viewed as a special case of fastENLOC with a special set of pre-defined enrichment parameters; the *coloc* algorithm converges to fastENLOC if the same enrichment parameters are supplied and the OCV assumption is satisfied.

### 6.2 Examining sensitivity of prior impact

To isolate the impact of prior specification on quantifications of colocalization, we consider a single SNP with various levels of evidence of associations from GWAS and eQTL analysis. In this experiment, we assume that priors for GWAS and eQTL associations are pre-determined at *p*_*d*_ = 10^−3^ and *p*_*γ*_ = 5 *×* 10^−5^, respectively. The values roughly reflect the prevalence of eQTL and GWAS hits in practice. Given *p*_*d*_ and *p*_*γ*_, the value of *α*_0_ is determined by *α*_1_ by

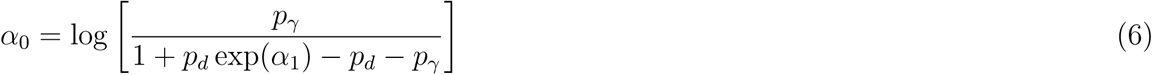

In this experiment, we consider only one independent genetic variant. To characterize the association evidence for each trait, we denote *q*_*d*_ and *q*_*γ*_ as the marginal posterior association probabilities of the target SNP for eQTL and GWAS, respectively. Subsequently, the marginal likelihood of associations, or Bayes factors, can be deduced by

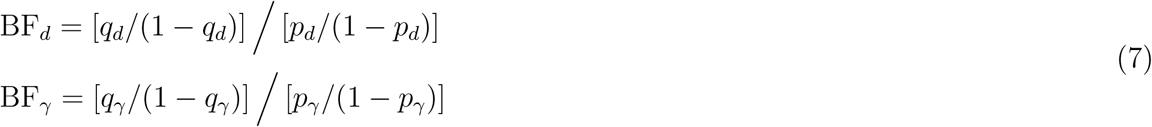

The posterior probabilities, *q*_*d*_ and *q*_*γ*_, are naturally scaled in [0, 1], and are convenient to classify different magnitudes of association evidence. (In comparison, because of the general difference in *p*_*d*_ and *p*_*γ*_, the direct comparison of Bayes factors is not as straightforward.) In this experiment, we consider the values of *q*_*d*_ and *q*_*γ*_ = 0.90, 0.50, 0.05 to represent strong, modest, and weak evidence for association, respectively.

With *p*_*d*_, *p*_*γ*_, *q*_*d*_, *q*_*γ*_, and *α*_1_ all fully specified, the SNP-level colocalization probability (SCP) can analytically computed (Section 2 of the Supplementary Material) To inspect the sensitivity of SCP with respect to the enrichment value, we vary *α*_1_ over a wide range covering all values typically seen in data, from -6 to 8, and compute the corresponding SCP values. Utilizing the equivalence of parameterization between ENLOC and *coloc*, we also provide the derivation of SCP as a function of coloc’s *p*_12_. The numerical result of SCP with respect to *p*_12_ is given in Supplementary Figure S3.

### 6.3 Simulation to inspect effect of allelic heterogeneity

To investigate the impact of allelic heterogeneity in colocalization analysis, we design a simulation scheme with real genotype data obtained from the GTEx whole blood data. To isolate the impacting factors and simplifying interpretations, we use the same genotype data to simulate gene expressions and complex traits. Thus, the LD patterns are perfectly matching between the two datasets. We use the genotypes from 400 individuals from the GTEx whole blood samples for this simulation. We create 4 sets of combined GWAS and eQTL datasets corresponding to the 4 different scenarios described in the main text. These scenarios enable us to investigate potential false positive and false negative findings by different AH assumptions. For each scenario, we simulate gene expressions for 1,000 genes and the corresponding complex trait data for 400 individuals using standard linear models. We particularly fix all gene expression effect sizes to 2.5, and the complex trait effect sizes to 1.5. The residual error variance is set to 1 for all simulations. Our consideration for this simulation setting is to ensure the generated association signals are strong, hence the subsequent colocalization analysis can be robust to prior specifications (based on what we observe from the previous simulations). We pool the 4 simulated scenarios into 2 different datasets. The first dataset consists of scenarios 1, 2, and 3, while the dataset consists of scenarios 2, 3, and 4. Thus, each dataset has exactly 3,000 genes for analysis. The two different datasets have different enrichment parameters, which allows us to examine the accuracy of the enrichment estimation procedure.

For fastENLOC analysis, we first run multi-SNP association analysis of simulated GWAS and eQTL data using DAP-G for all simulated genes. For *coloc*, we directly supply summary statistics from the corresponding single-SNP analysis.

### 6.4 Simulation to benchmark power of colocalization analysis

To benchmark and investigate the performance of probabilistic colocalization analysis, we simulate complex trait and gene expression data that resemble observed data in practice. For this experiment, we use the real genotype data across 20,000 genes from 400 participants of the GTEx project. To ensure the LD patterns in genotypes are perfectly matched in eQTL and GWAS data, we again use the identical genotype data to independently simulate expression and complex traits.

For each candidate gene, we select a fixed number of 1,500 *cis*-SNPs whose minor allele frequencies > 0.03. We use the following linear model to simulate the expression level of gene *i* for individual *k*,

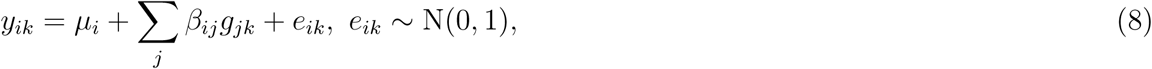

where *g*_*jk*_ and *β*_*ij*_ represent the genotype and the genetic effect for SNP *j* (*j* = 1, 2, *…*, 1500). Particularly, *β*_*ij*_ is independently drawn from a mixture distribution,

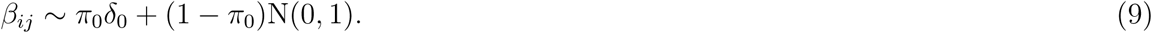

That is, with probability *π*_0_ = 1 − 3*/*1500 = 0.998, the genetic effect of SNP *j* on expression is exactly 0; and with probability 0.002, the SNP has a non-zero random effect drawn from *rmN* (0, 1) distribution. On average, the simulation scheme yields 3 causal eQTL SNPs per *cis* region.

To simulate the causal GWAS hits, we conduct an independent Bernoulli trial on each candidate SNP: if a SNP is not an eQTL SNP, its probability of being a causal GWAS SNP is set to 1/1500; otherwise, the corresponding probability increases to 0.035, or equivalently, *α*_1_ = 4 (which is similar to the enrichment level of blood eQTLs in causal GWAS hits of lipids traits [14]). Subsequently, we again simulate the complex trait using a multiple linear regression model. Specifically, the genetic effect of each causal GWAS SNP is drawn from the distribution N(0, 1) and the residual error is also simulated from the same distribution.

Overall, across 20,000 genes, this scheme generates 59,937 causal eQTL SNPs and 22,054 casual GWAS hits. There are 2,103 instances that the causal variants for both traits are overlapped.

### 6.5 Simulation to investigate effect of LD mismatch

We construct these simulations based on the design of the GEUVADIS project, which studies the eQTLs across 5 different populations (FIN, CEU, GBR, TSI, and YRI). We use the real genotype data for the five populations that are originally genotyped in the 1000 Genome project Phase I [26]. We select 6,977 genes from the GEUVADIS project that contain at least 500 SNPs in a 200 KB *cis*-region. Note that we keep all SNPs in all studied populations, even though some of them are monomorphic or extremely rare in specific population groups. We apply a similar scheme as the previous simulation (Section 6.4) to sample the causal eQTL and GWAS SNPs based on independent Bernoulli trials and pre-specified enrichment parameters. We adjust the enrichment parameter *α*_1_ from 4.0 to 4.5 to increase the instances of colocalizations and compensate reduced numbers of *cis*-SNPs.

Given the causal eQTL and GWAS status, we simulate a single set of eQTL data using the genotypes from the FIN population. Next, we simulate 5 different sets of complex trait data for all populations. The sample sizes for each population are 92 (FIN), 81 (CEU), 87 (GBR), 95 (TSI), and 80 (YRI), respectively. The phenotype simulation is based on multiple linear regression models, and the residual error variance for each corresponding linear model is also set to 1. To compensate for the reduced sample size, we draw the genetic effects for both eQTLs and GWAS hits from a random effect model with variance = 2. In total, the simulated datasets contain 20,987 eQTLs and 9,378 GWAS hits with 2,488 colocalized signals.

### 6.6 Colocalization analysis of 4,091 complex traits with GTEx eQTL data

The eQTL data from 49 tissues are generated from the final version of the GTEx project. The eQTL data are processed based on the protocols established by the GTEx consortium. The processing pipeline and related software packages are provided in [18]. We collect the summary statistics in the form of single-SNP testing *z*-scores from 4,091 GWAS of complex traits. These summary statistics are harmonized to be compatible to be analyzed with the GTEx data. In particular, additional *z*-scores are imputed according to LD patterns observed in the GTEx data using the software package LDpred [32]. The technical details on the pre-processing of the complex data are also documented in [18, 6].

The multi-SNP fine-mapping analysis of the eQTL in each tissue is performed using DAP-G, and the resulting probabilistic annotations of eQTLs are available in GTEx portal. Due to a lack of individual-level data or precise LD information for the complex trait GWAS, we choose to employ the fine-mapping algorithm similar to *fgwas* implemented in the software package TORUS [16]. Briefly, TORUS segments the genome into 1 to 2 Mb wide LD blocks [33], and performs fine-mapping analysis within each LD block assuming a single causal GWAS hit. Although the assumption is imperfect, we are much less likely to observe multiple strong GWAS hits within a single LD block in practice.

We estimate the enrichment levels of eQTLs in the GWAS hits for each tissue-trait pair using fastENLOC. Particularly, we experiment two different strategies for estimation: with and without applying shrinkage to enrichment estimates. Our observation is that in many cases where strong evidence for colocalization events is lacking, the direct estimates of *α*_1_ without shrinkage are often unstable. That is, the point estimates can be wildly positive or negative and they are always associated with substantially large variances. The corresponding *z* scores, which measure the signal-to-noise ratio, are almost never statistically significant. In these scenarios, direct use of the point estimates while ignoring the large standard errors is potentially dangerous for downstream colocalization analysis. Hence, we conclude it is necessary to apply shrinkage to the estimate of *α*_1_. Although it is known that the shrinkage estimates are biased toward 0, the tradeoff to stabilize the estimates is necessary in this context. The implementation of fastENLOC assumes a normal prior, N(0, 1*/λ*) prior on *α*_1_, which has the ability to induce a shrinkage point estimate. The prior precision parameter *λ* defines the level of shrinkage, and *λ* is set to 1 by default. This prior shrinkage effectively shrinks the unstable estimates to near 0 but has little impacts on the estimates that are originally stable. The histograms contrasting the estimates with and without shrinkage are shown in Supplementary Figure S2.

