## Supplementary Materials for "Probabilistic Colocalization of Genetic Variants from Complex and Molecular Traits: Promise and Limitations"

### 1 Constraints in ENLOC parameterization

Here we show that the combination of the three prior parameters, $(p_\gamma, p_d, \alpha_1)$, fully specifies the prior model for SNP-level colocalization analysis. Note that ENLOC/fastENLOC uses an alternative parameterization with $(p_\gamma, \alpha_0, \alpha_1)$. It is therefore sufficient to show that $\alpha_0$ can be expressed using $(p_\gamma, p_d, \alpha_1)$.

Note that

$$
\begin{aligned}
p_\gamma &= \Pr(\gamma = 1 \mid d = 0)\,(1 - p_d) + \Pr(\gamma = 1 \mid d = 1)\,p_d \\
&= \frac{\exp(\alpha_0)}{1 + \exp(\alpha_0)}\,(1 - p_d) + \frac{\exp(\alpha_0 + \alpha_1)}{1 + \exp(\alpha_0 + \alpha_1)}\,p_d
\end{aligned} \tag{1}
$$

Thus, $\alpha_0$ can be solved from the above equation given $p_\gamma, p_d$, and $\alpha_1$. Furthermore, an accurate approximation of $\alpha_0$ can be analytically computed by noting that, in practice, $\Pr(\gamma = 1 \mid d = 0) \ll 1$ and $\Pr(\gamma = 1 \mid d = 1) \ll 1$ (i.e., GWAS hits for complex traits are sparse). It follows that

$$
(1 + \exp(\alpha_0))\; p_\gamma \approx \exp(\alpha_0)\,(1 - p_d) + \exp(\alpha_0 + \alpha_1)\,p_d \tag{2}
$$

and

$$
\alpha_0 \approx \log\left(\frac{p_\gamma}{1 + p_d \exp(\alpha_1) - p_d - p_\gamma}\right) \tag{3}
$$

### 2 Computing SNP-level Colocalization Probability

Here we derive the SNP-level colocalization probability (SCP) for an independent SNP given the posterior association probabilities, $q_d := \Pr(d = 1 \mid \text{eQTL data})$ and $q_\gamma = \Pr(\gamma = 1 \mid \text{GWAS data})$ for association analysis of individual traits.

By the Bayes rule, the Bayes factor for eQTL association can be computed by

$$\begin{aligned} \mathrm{BF}_d &= \frac{\Pr(\text{eQTL data} \mid d = 1)}{\Pr(\text{eQTL data} \mid d = 0)} \\ &= \left[q_d/(1-q_d)\right] \Big/ \left[p_d/(1-p_d)\right]. \end{aligned}$$

Similarly, the Bayes factor for GWAS association at the target SNP is given by

$$\mathrm{BF}_\gamma = \left[q_\gamma/(1-q_\gamma)\right] \Big/ \left[p_\gamma/(1-p_\gamma)\right].$$

Assuming a two-sample design, it follows that

$$\begin{aligned} &\Pr(\text{GWAS data, eQTL data} \mid d, \gamma) \\ &= \Pr(\text{eQTL data} \mid d)\ \Pr(\text{GWAS data} \mid \gamma) \end{aligned} \tag{4}$$

Applying the Bayes rule to compute SCP yields

$$\begin{aligned} &\Pr(d = 1, \gamma = 1 \mid \text{eQTL data, GWAS data}) \\ &= \frac{\Pr(d=1,\gamma=1)\mathrm{BF}_d\mathrm{BF}_\gamma}{\Pr(d=0,\gamma=0) + \Pr(d=1,\gamma=0)\mathrm{BF}_d + \Pr(d=0,\gamma=1)\mathrm{BF}_\gamma + \Pr(d=1,\gamma=1)\mathrm{BF}_d\mathrm{BF}_\gamma}, \end{aligned} \tag{5}$$

where

$$\begin{aligned} \Pr(d=1,\gamma=1) &= \frac{\exp(\alpha_0+\alpha_1)}{1+\exp(\alpha_0+\alpha_1)}\, p_d \\ \Pr(d=0,\gamma=0) &= \frac{1}{1+\exp(\alpha_0)}\,(1-p_d) \\ \Pr(d=1,\gamma=0) &= \frac{1}{1+\exp(\alpha_0+\alpha_1)}\, p_d \\ \Pr(d=0,\gamma=1) &= \frac{\exp(\alpha_0)}{1+\exp(\alpha_0+\alpha_1)}\,(1-p_d) \end{aligned} \tag{6}$$

### 3 Calibration of Regional Colocalization Probabilities

In the simulations of assessing empirical power of colocalization analysis, we also examine the calibration of the regional colocalization probabilities (RCPs) reported by both fastENLOC and *coloc*. The concept of calibration refers to where reported Bayesian posterior probabilities represent the frequencies of events in multiple independent experiments, a frequentist property. To this end, we sort all RCP values into evenly-divided probability bins (e.g., $[0, 0.1), [0.1, 0.2), ..., [0.9, 1.0]$). Within each probability bin, we then compute the fractions of reported sites that truly harbor a colocalized signal. If the RCP values are calibrated, we expect that the mean RCP from each pre-defined bin aligns with the corresponding fraction of true colocalized signals. Overall, we find that the calibration of the computed RCPs are overly conservative (Figure S1). Applying the true enrichment parameters improves the overall calibration but does not completely resolve the issue of conservativeness, especially for the RCPs in the low to modest range. This is also in agreement with our main conclusions from the power and type I errors of the colocalization analysis, in Table 2 of the main text. We also conduct a similar analysis for the gene-level posterior probability of colocalization from the *coloc* analysis, which confirms that inaccurate prior information and/or ignoring potential AH could lead to anti-conservative assessment of colocalization probabilities.

### 4 Shrinkage estimation of enrichment parameter

Let $\tilde{\alpha}_1$ and $v_1$ denote the point estimate of $\alpha_1$ and the corresponding variance directly obtained from the multiple imputation procedure of ENLOC. In the case of a lack of strong evidence of colocalization events, $\tilde{\alpha}_1$ can be highly unstable, i.e., $\tilde{\alpha}_1$ can take extreme positive or negative values, and $v_1$ is also extremely large. The phenomenon is similar to the enrichment estimation using a $2 \times 2$ contingency table where at least one cell count $\to 0$. To stabilize the estimate in the empirical Bayes framework, we consider a shrinkage prior $\mathrm{N}(0, 1/\lambda)$ for $\alpha_1$. The resulting shrinkage estimate of the $\alpha_1$ is given by

$$\hat{\alpha}_1 = \frac{\tilde{\alpha}_1}{1 + \lambda v_1}, \tag{7}$$

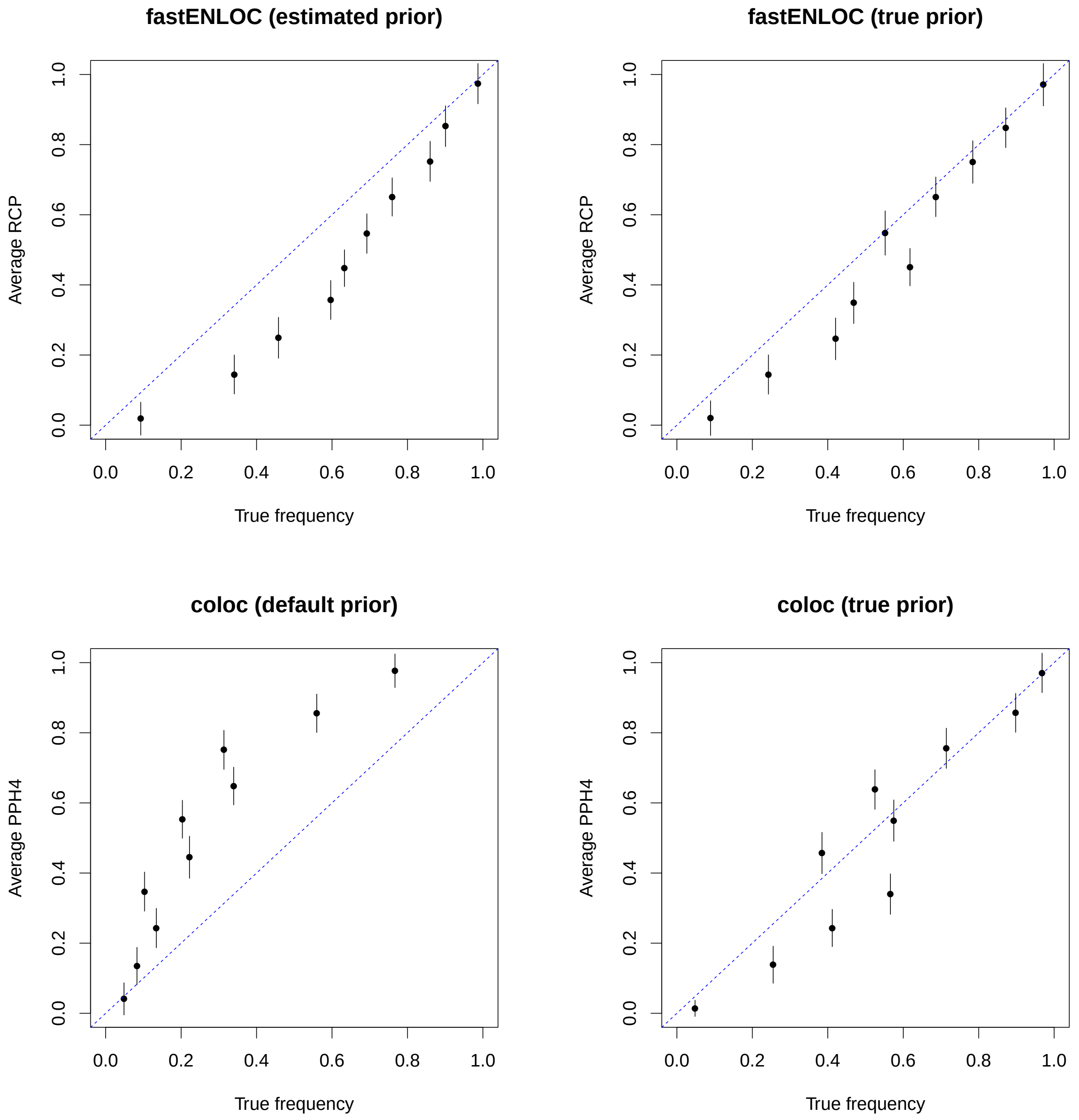


Figure S1: **Calibration of reported regional colocalization probabilities** The error bars represent 95% confidence intervals. The RCPs from the fastENLOC are conservative in comparison to the corresponding frequencies. The conservativeness is partially explained by the conservative estimation of the enrichment priors. In the case of *coloc*, the default prior leads to severely anti-conservative results. Even when the true priors are applied, the *coloc* results still show anti-conservativeness in some frequency bins.

and the corresponding variance is

$$\text{Var}(\hat{\alpha}_1) = \frac{v_1}{1 + \lambda v_1} \tag{8}$$

The shrinkage parameter $\lambda_1$ defines the strength of the shrinkage. As $\lambda_1 \to 0$, it follows that $\hat{\alpha}_1 \to \tilde{\alpha}_1$ and $\text{Var}(\hat{\alpha}_1) \to v_1$, i.e., there is no effect of shrinkage. On the other extreme, as $\lambda \to \infty$, it follows that $\hat{\alpha}_1 \to 0$ and $\text{Var}(\hat{\alpha}_1) \to 0$. The default implementation of fastENLOC set $\lambda = 1$.

We observe that the shrinkage estimates of the enrichment parameter are effectively stabilized. The comparison of $\tilde{\alpha}_1$ and $\hat{\alpha}_1$ from analyzing 4,091 complex traits and GTEx eQTL data are shown in Figure S2

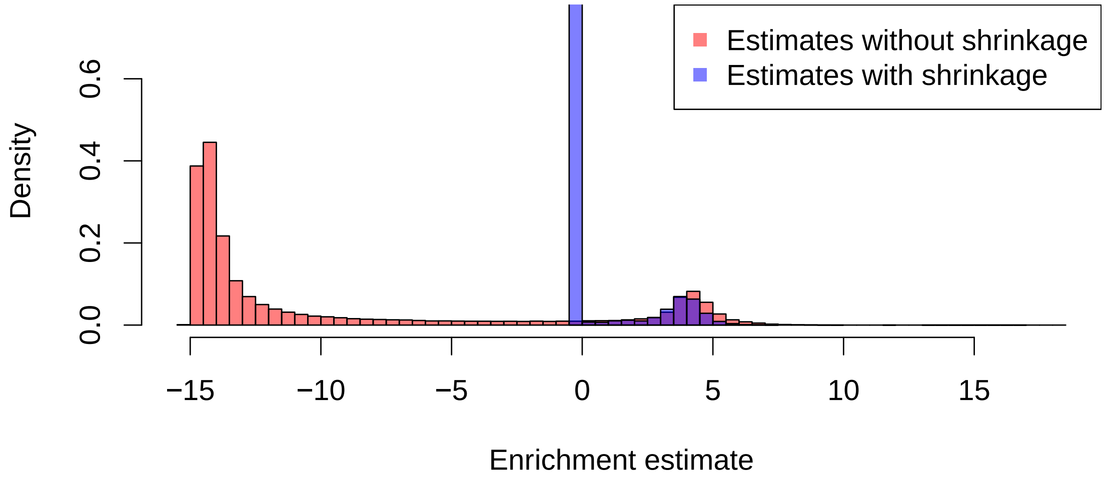


Figure S2: **Enrichment estimates with and without shrinkage from all tissue-trait pairs in the phenomexcan analysis**

### 5 Additional Supplementary Figures

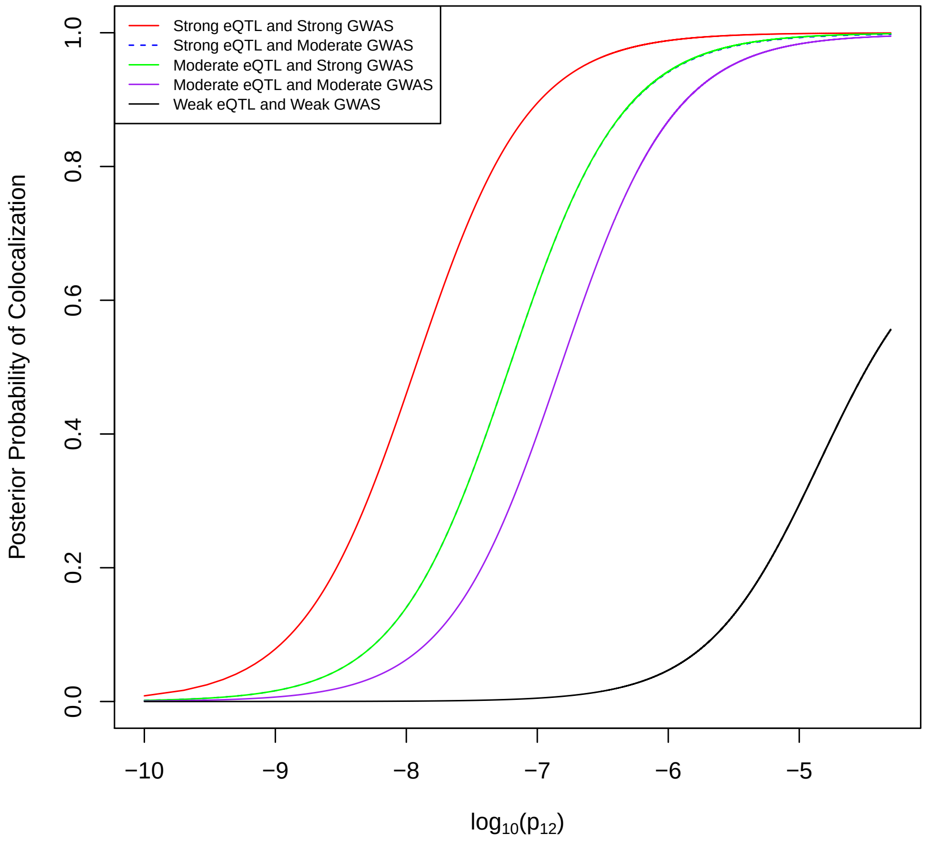


Figure S3: **An illustration of the sensitivity of SNP-level colocalization probability with respect to *coloc* parameterization of $p_{12}$** The range of the values for $p_{12}$ correspond to the $\alpha_1$ values in Figure 1 of the main text.
